# Distinct profiles of LRRK2 activation and Rab GTPase phosphorylation in clinical samples from different PD cohorts

**DOI:** 10.1101/2021.11.05.465894

**Authors:** Lilian Petropoulou-Vathi, Athina Simitsi, Politymi-Eleni Valkimadi, Maria Kedariti, Lampros Dimitrakopoulos, Christos Koros, Dimitra Papadimitriou, Alexandros Papadimitriou, Leonidas Stefanis, Roy N. Alcalay, Hardy J. Rideout

## Abstract

Despite several advances in the field, pharmacodynamic outcome measures reflective of LRRK2 kinase activity in clinical biofluids remain urgently needed. A variety of targets and approaches have been utilized including assessments of LRRK2 itself (levels, phosphorylation), or its substrates (e.g. Rab10 or other Rab GTPases). We have previously shown that intrinsic kinase activity of LRRK2 isolated from PBMCs of G2019S carriers is elevated, irrespective of disease status. In the present study we find that phosphorylation of Rab10 is also elevated in G2019S carriers, but only those with PD. Additionally, phosphorylation of this substrate is also elevated in 2 separate idiopathic PD cohorts, but not in carriers of the A53T mutation in α-synuclein. In contrast, Rab29 phosphorylation was specifically reduced in urinary exosomes from A53T and idiopathic PD patients. Taken together, our findings highlight the need for the assessment of multiple complimentary targets for a more comprehensive picture of the disease.

---

The kinase activity of leucine-rich repeat kinase 2 (LRRK2) has emerged as the primary target for both therapeutic development and biomarker design. Both cellular and animal models have demonstrated that neurodegeneration induced by mutant forms of LRRK2 is kinase activity-dependent (e.g. ^1^). Similarly, early biomarker studies, assessing LRRK2 auto-phosphorylation (pS1292-LRRK2) in urinary exosomes, have indicated that LRRK2 activity may be elevated during the clinical progression of LRRK2-associated, as well as idiopathic, PD ^2,3^. In addition to auto-phosphorylation of LRRK2 and its intrinsic kinase activity, which are measures of distinct aspects of its kinase function, other readouts of the activation state of LRRK2 include phosphorylation of endogenous substrates, such as members of the Rab family of small GTPases ^4^. Extending these findings, the group of Greenamyre and colleagues ^5^ applied a novel proximity ligation assay to detect pS1292-LRRK2 and pT73-Rab10 in ventral midbrain of post mortem tissue from iPD patients. In subsequent studies, we have reported that the *in vitro* kinase activity of purified LRRK2 from PBMCs of G1029S-LRRK2 mutation carriers is elevated ^6^. Alternatively, while phosphorylation at the S935 site within the N-terminal region of LRRK2 is a robust measure of target engagement (e.g. ^7,8^), it does not completely track with kinase activity of LRRK2 ^9^.

Rab10 phosphorylation has been assessed in various PD cohorts, however the findings have been mixed. In a small proof-of-concept study, Rab10 phosphorylation (T73) in neutrophils was not significantly changed in carriers of the G2019S LRRK2 mutation ^10^; and in PBMCs from iPD patients, pT73-Rab10 was not changed compared to healthy controls, despite being correlated with levels of certain cytokines ^11^. Interestingly, in neutrophils from carriers of the less common R1441G mutation in LRRK2, a significant increase in pT73-Rab10 is detected in comparison to healthy controls, irrespective of disease status ^12^. This is consistent with findings reported in transgenic mice expressing R1441C or G2019S, where a robust increase in Rab10 phosphorylation was observed in R14411C but not G2019S mice ^13^. Most recently, using a proprietary antibody raised against pT73-Rab10, Wang and colleagues reported that phosphorylated Rab10 levels, normalized to total LRRK2, are increased in PBMCs of affected carriers of the G2019S mutation ^8^ and are reduced in cells treated *ex vivo* with LRRK2 kinase inhibitors.

In the present study, we find increased Rab10 phosphorylation in iPD patients and *affected* G2019S-*LRRK2* carriers only, whereas levels in healthy carriers and A53T-*SNCA* PD patients are not changed from healthy controls. Additionally, phosphorylation of Rab29 is decreased in urinary exosomes from *SNCA* mutation carriers and iPD patients. Thus, it appears that alterations in peripheral LRRK2-dependent Rab phosphorylation are not uniformly linked to either disease status or genotype.

## Study participants

The samples that we assessed were collected as part of two separate biomarker studies. The demographics of a subset of the clinical cohort (healthy controls, iPD, G2019S-*LRRK2*/PD+ & G2019S-*LRRK2*/PD-) followed at Columbia University Medical Center (New York, USA) have already been reported in a separate manuscript ^6^. The second cohort (healthy controls, iPD, & A53T-*SNCA* PD) was derived from patients followed at the Departments of Neurology at Henry Dunant Hospital Center (Athens, Greece) and Aiginiteio University Hospital (Athens, Greece). Thus, in this study we have included two independent groups of healthy controls and iPD patients. Demographics and key clinical features of the Athens cohort are summarized in Supplementary Table 1. The samples from each cohort were analyzed independently. In some cases, fewer samples were included in a specific analysis than were originally collected due to technical issues with detection of the specific target in the sample matrix. For example, in the samples where signal from the total protein (for normalization) was missing, these were excluded from the analyses.

## PBMC & Urinary exosome isolation

PBMCs were isolated from whole blood collected in Heparin-coated Vacutainer collection tubes (BD), using protocols established in our recent work ^6^. Some aliquots were divided into 2 samples and treated with DMSO or the LRRK2 inhibitor MLi-2, then collected/washed in PBS and snap frozen as a cell pellet in dry ice, and stored at − 80°C. Urine (~50-200 ml) was collected from participants and tested for urinary infections. Exosomes were isolated as described from the group of West and colleagues ^3^, and lysed under the same conditions as PBMCs. Extracts of PBMCs or urine EVs were subjected to SDS-PAGE and probed for total and phosphorylated (T73) Rab10, and total or phosphorylated (T71) Rab29. For quantification and normalization of the phosphorylated protein, the ratio of band intensity of phosphorylated:total protein was determined. When possible, an internal control, comprised of a parallel sample, was included on each gel; otherwise, band intensities were measured from films taken from equivalent exposures.

## Statistical Analyses

For statistical comparison of the different study groups, we employed Mann-Whitney U non-parametric tests & Bonferroni’s post-hoc test; ANOVA, with Tukey’s post-hoc multiple comparisons; and two-tailed t-tests.

## Phosphorylation of Rab10, but not Rab29, is increased in specific PD cohorts

By Western immunoblotting, we assessed levels of pT73-Rab10 and pT71-Rab29 in PBMC extracts from each group. In Figure 1a, a representative immunoblot showing levels of phosphorylated (T73) and total Rab10 from each group is shown; with quantification of the bands presented in Figure 1b. When normalized to total Rab10 levels, we found a significant increase in pT73-Rab10 in idiopathic PD (iPD) from both the Columbia cohort, as well as the Athens cohort, compared to healthy control subjects (Fig. 1 a,b). Like LRRK2 expression ^6^, expression of Rab10 (total) was not significantly altered in either iPD cohort.

**Figure 1.**
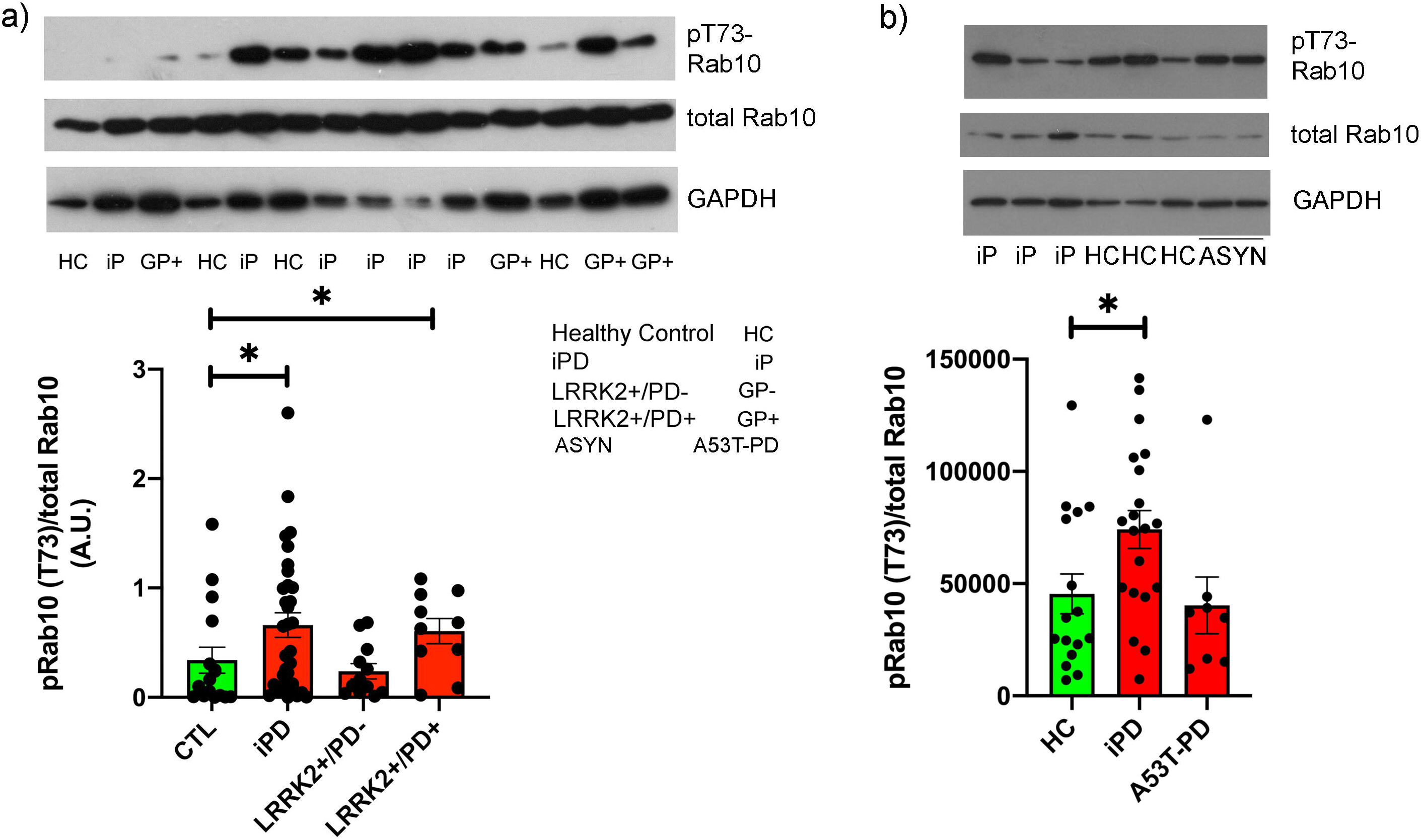
Phosphorylation of Rab10 is elevated in PBMCs from idiopathic PD and affected carriers of the G2019S mutation in LRRK2. Extracts of PBMCs from the indicated subject groups were separated by SDS-PAGE and probed for phosphorylated (T73) and total Rab10, as well as GAPDH. (A) Representative western immunoblots showing pT73-Rab10 and total Rab10; with the quantification of the band intensity shown below. Mann-Whitney tests were performed to identify significant differences between groups. * p<0.05. (B) PBMC extracts were separated by SDS-PAGE. A representative western immunoblot probed for pT73-Rab10 and total Rab10; with the band intensities quantified and presented below. For normalization, total expression levels of Rab10 were normalized to GAPDH first, and the band intensity of pT73-Rab10 was then normalized to the corrected total Rab10. Mann-Whitney tests were performed to identify significant differences between groups. * p<0.05.

In the two G2019S-*LRRK2* carrier groups, we found a divergence in Rab10 phosphorylation that was strictly linked to disease status. Affected carriers of this mutation exhibited a significant increase in normalized pT73-Rab10 levels, similar to iPD patients, compared to healthy controls; whereas pRab10 levels were not different from control subjects in non-manifesting G2019S-*LRRK2* carriers (Fig. 1 a,b). We have previously shown from the same cohort ^6^, that the *in vitro* kinase activity of LRRK2 purified from PBMCs of both G2019S carrier groups is significantly increased compared to healthy control and iPD patients. Thus, in a cellular milieu, the disease context is clearly impacting the phosphorylation of LRRK2 substrates, even in the presence of a hyperactive enzyme. Finally, in PBMCs from affected carriers of the A53T mutation in α-synuclein, phosphorylation of Rab10 was unchanged in PBMCs compared to healthy controls (Fig. 2 a,b).

**Figure 2.**
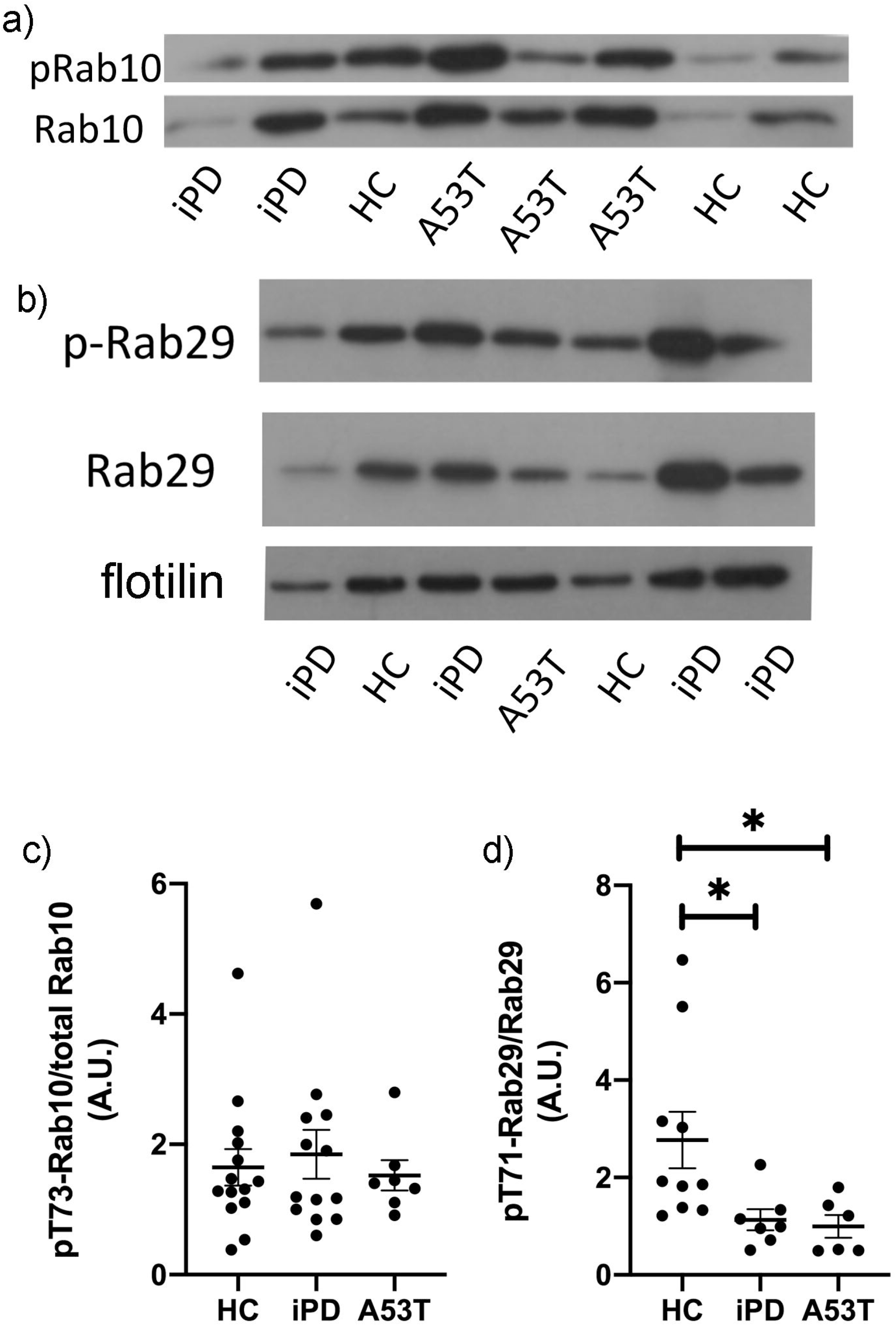
Phosphorylation of Rab10 is unaffected in urinary exosomes but phosphorylation of Rab29 is decreased in two PD cohorts. Exosomes from urine were purified by ultracentrifugation, and extracts separated by SDS-PAGE. The membranes were probed with total or phosphorylated Rab10 and 29, and flotilin as a marker for exosomes. Band intensities were normalized to flotilin and the ratio of phosphorylated Rab to total Rab. (A & B) Representative blots showing phosphorylated Rab10 and Rab29; and quantified in (C & D). Mann-Whitney tests were performed to identify significant differences between groups. * p<0.05

In addition to Rab10 phosphorylation, we assessed the induction of phosphorylation of another LRRK2 substrate, Rab29, in PBMCs from each group. We could readily detect robust expression of Rab29 in PBMCs from each group, and this did not significantly differ between each group. Interestingly, in most samples, we could not detect phosphorylated Rab29 (T71), and when it was detected, it was possible only in the presence of a signal boost reagent (SignalBoost Immunoreaction, Sigma; not shown), suggesting that within PBMCs, Rab29 is not a particularly prominent LRRK2 substrate, either in the healthy control population or in any of the PD cohorts; or that specific Rab29 phosphatases are particularly active in these cells.

## Changes in Rab10 and Rab29 phosphorylation in urinary exosomes

In extracts of urinary exosomes, the detection of LRRK2 was highly variable; in many samples, total and pS1292-LRRK2 was un-detectable (Supplemental Figure 1b), even in the presence of the SignalBoost detection reagent. However, we were able to easily detect both total and pT73-Rab10 in extracts of urinary EVs by Western immunoblotting (Fig. 2a). The pattern of Rab10 phosphorylation differed in urinary EVs; where, in contrast to PBMCs, phosphorylated Rab10, normalized to total Rab10, in EVs from iPD and A53T-αsyn patients were unchanged compared to healthy controls (Fig. 2b). Interestingly, whereas we could not detect phosphorylated Rab29 in extracts of PBMCs from this cohort, we were able to detect pT71-Rab29 in urinary EVs; and found a significant decrease in pRab29 (normalized to total Rab29) in EVs from A53T-PD patients compared to healthy controls (Fig. 2c). Phosphorylated Rab29 levels in urinary EVs from iPD patients were also reduced compared to healthy controls (Fig. 2c).

In the present study, we assessed Rab GTPase phosphorylation in PBMCs and urinary exosomes from multiple PD cohorts collected at different clinical centres. We recently reported several LRRK2-based outcome measures in a cohort of G2019S-*LRRK2* carriers (both healthy and affected) from the CUMC-Movement Disorders centre. In PBMCs from these individuals, we found a significant elevation in phosphorylation of the LRRKtide peptide substrate in both G2019S carrier groups, compared to healthy controls and idiopathic PD patients ^6^. This is not surprising given the conserved motif that this mutation lies within in the LRRK2 kinase domain (e.g. ^14^). In the same report, we could not detect changes in the *in vitro* kinase activity of LRRK2 isolated from iPD PBMCs ^6^.

In PBMCs from G2019S carriers, we report here increased phosphorylation of Rab10 (T73) *only* in affected carriers of this mutation, healthy G2019S carriers exhibited pT73-Rab10 levels similar to healthy controls. Thus, the isolated enzyme bearing the Gly to Ser substitution in the critical *DYG* kinase domain motif exhibits elevated intrinsic kinase activity, regardless of disease status ^6^; however, in a cellular context the disease status clearly interacts with the genotype to modulate Rab10 phosphorylation, at least in peripheral immune cells. In other words, *in vitro* kinase activity is a good predictor of LRRK2 mutation status, but not of disease status. In another recent study, combining western immunoblotting and targeted mass-spectrometry measures of phosphorylated Rab10, it was reported that T73-Rab10 levels in a small sample of carriers of the R1441G were significantly elevated in neutrophils compared to healthy control subjects ^12^. Unlike the present study, phosphorylated Rab10 levels from carriers of the G2019S-*LRRK2* mutation were not different from healthy controls, regardless of disease status ^12^; however in an earlier study also using an MS-based approach, pT73-Rab10 levels in neutrophils were approximately 2-fold higher in a set of G2019S carriers ^15^.

In PBMCs from iPD or A53T-PD patients, we do not detect changes in the intrinsic kinase activity of isolated LRRK2 (^6^; and manuscript in preparation); whereas in two independent cohorts (CUMC and Athens), we find increased phosphorylation of Rab10 in PBMCs from iPD patients. Previous reports have not revealed differences in Rab10 phosphorylation in this patient group (e.g. ^11^); the reasons for this discrepancy are unclear, however may be related to the normalization methods taken in each study.

A second key finding we report here is the loss of phosphorylation of Rab29 (Thr71) in urinary exosomes from iPD and A53T-PD patients. Recently, it was reported by two independent groups that over-expression of Rab29 can trigger the recruitment of LRRK2 to intracellular membranes and the trans-Golgi network (TGN) leading to its activation ^16,17^; suggesting a regulatory cycle between LRRK2 activity and Rab29. In our clinical biofluid samples, the *loss* of phosphorylation of Rab29 in urine exosomes from iPD and A53T-PD patients, releasing its inhibition, would be consistent with increased activation of LRRK2 (e.g. increased pT73-Rab10 in the iPD group). While we could not detect changes in phosphorylation of Rab10 in this cohort, the reduction in Rab29 phosphorylation is the first indication of possible changes in LRRK2 activation in clinical samples from PD patients carrying the A53T-*SNCA* mutation; and bolsters the notion that LRRK2 function plays an important role in PD in general. A summary of our findings in the current study, in the context of our previously reported findings from one of the two cohorts (from CUMC) is presented in Table 2. Our findings reinforce the need to expand our pallet of validated LRRK2-pathway outcome measures to include additional targets as well as multiple biofluid types ^18^.

**Table 1.** Summary of demographics of study participants (Athens cohort).

|  | **Healthy Controls (HC)** | **Idiopathic PD (iPD)** | **A53T-SNCA PD** | **p-value** | **Statistical tests** |
| --- | --- | --- | --- | --- | --- |
| **Gender (M:F)** | 7:9 | 14:6 | 4:4 |  |  |
| **Age** | 55 (36-75) | 72 (47-85) | 51 (30-73) |  |  |
| **Age at onset** | na | 66 (42-80) | 46 (27-68) | p=0.0011 | MWU (iPD vs A53T-PD) |
| **LEDD** | na | 629 (0-1865) | 758 (150-1500) | ns | MWU (iPD vs A53T-PD) |
| **UPDRS-III** | 0 | 30 (0-91) | 31 (9-53) | ns | MWU (iPD vs A53T-PD) |
| **MoCA** | 26 (17-30) | 19 (4-26) | 26 (16-30) | HC vs. iPD (p=0.0025)<br>HC vs. A53T (ns) | MWU |
| **H&Y** | 0 | 2 (0-4) | 2 (1-3) | ns | MWU (iPD vs A53T-PD) |

**Table 2.** Summary of LRRK2-specific changes in two PD cohorts.

|  | **Total LRRK2** | ***In vitro* kinase activity** | **pS935-LRRK2** | **pS1292-LRRK2 (WB)** | **pT73-Rab10 (WB)** | **pT71-Rab29 (WB)** |
| --- | --- | --- | --- | --- | --- | --- |
| **Healthy Control (HC)** | Baseline | Baseline | Baseline | Detectable only | Baseline | Baseline |
| **Idiopathic PD (iPD)** | Like HC | Like HC | Elevated (2 cohorts; Melachroinou et al., 2020; manuscript in prep.) | Detectable only | Elevated (2 cohorts) | Decreased (Ur-EVs) |
| **G2019S+ PD+** | Like HC | Elevated (Melachroinou et al., 2020) | Slight decrease (Melachroinou et al., 2020) | Detectable only | Elevated | ND |
| **G2019S+ PD-** | Like HC | Elevated (Melachroinou et al., 2020) | Slight decrease (Melachroinou et al., 2020) | Detectable only | Like HC | ND |
| **A53T-SNCA PD** | Like HC | Like HC | Like HC | Detectable only | Like HC | Decreased (Ur-EVs) |

---

**Supplemental Figure 1**. **LRRK2-dependent phosphorylation of Rab10 in PBMCs and the detection of pS1292-LRRK2.** (A) PBMCs from healthy control individuals were treated with vehicle or the indicated concentrations of MLi-2 for 2hr; the cell extracts were then separated by SDS-PAGE, and the membranes probed for total and phosphorylated (T73) Rab10 and β-actin. In panel (B), a representative immunoblot is shown following probing membranes for total and phosphorylated pS1292-LRRK2 in the presence of the SignalBooster Immunodetection reagent. (C) An aliquot of urine exosomes resuspended in PBS was removed and prepared for TEM imaging. A representative micrograph is shown depicting several exosomes (red arrow). Scale bar = 200nm.

