## Supplementary figures and images for "Distinct profiles of LRRK2 activation and Rab GTPase phosphorylation in clinical samples from different PD cohorts"

### Supplemental Data

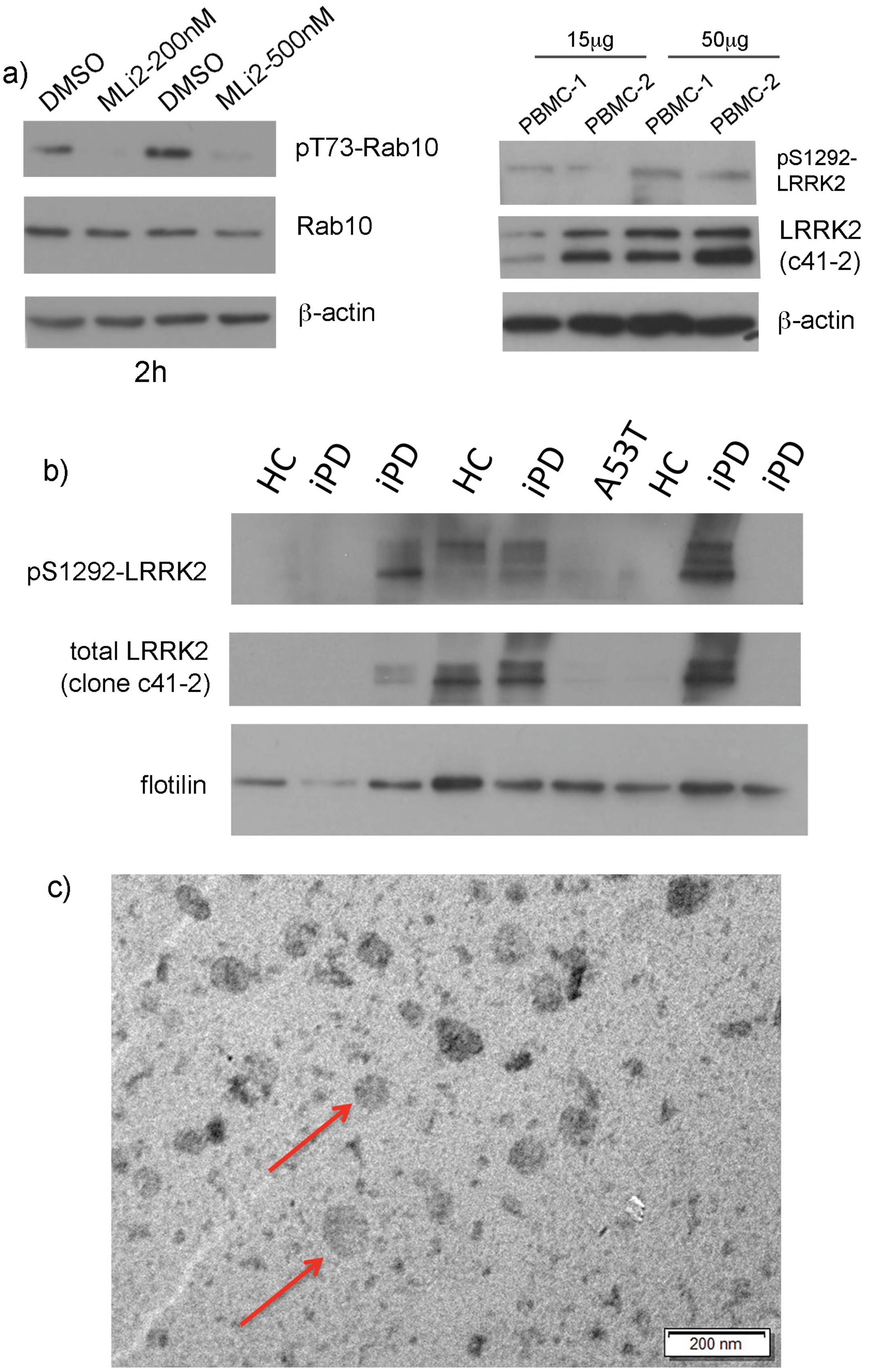


Supplemental Fig. 1; Petropoulou-Vathi et al.
