## Supplementary Material for "Distinct profiles of LRRK2 activation and Rab GTPase phosphorylation in clinical samples from different PD cohorts"

**Materials & Methods**.

After the final wash, PBMC pellets were lysed as described in ^1^. For western immunoblotting, protein samples were separated by SDS-PAGE, and the membranes blocked in either 5% Milk/TBST (for Rab10, Rab29, GAPDH, LRRK2, and Flotilin-1) or in 5% BSA/TBST (for pS1292-LRRK2, pT73-Rab10 and pT71-Rab29), for 1h at RT. The primary antibodies used were as follows: Rab10 (ab237703 Abcam, rabbit, 1:1000), Rab29 (ab199644 Abcam, rabbit, 1:1000), Flotilin-1 (sc-133153 Santa Cruz, mouse, 1:1000), LRRK2 (ab133474 Abcam rabbit, clone c41-2, 1:1000), pT73-Rab10 (ab230261 Abcam, rabbit, 1:1000) and pT71 Rab29 (ab241062 Abcam, rabbit, 1:1000), and pS1292-LRRK2 (ab203181 Abcam, clone MJF-19-7-8, 1:1000), shaking overnight at 4^o^C. In some cases, where the membranes were probed with anti pS1292-LRRK2 antibodies, we included Signal Boost Immunoreaction Reagent (Sigma) in the antibody diluent for both the primary and secondary antibody incubation steps. Membranes were then washed 3x for 10min in 1X TBST and incubated with HRP-conjugated secondary antibodies (Millipore AP132P anti-rabbit or Millipore AP124P anti-mouse, 1:5000), for 1h at RT. Three more washes followed and membranes were exposed to ECL and developed using an automatic film developer. Films were scanned and quantified using ImageJ, with the final Figures compiled using Adobe Photoshop.

For the preparation of urinary exosomes for visualization by transmission electron microscopy (TEM), the exosome pellets were resuspended in PBS, and 2 μl of exosome suspension removed for imaging. The samples were processed as described ^2^.

**Results**.

*Phosphorylated S1292-LRRK2 is barely detectable in peripheral blood cells of PD patients*. Prior to assessments of Rab phosphorylation in clinical samples as an index of LRRK2 kinase activity, auto-phosphorylation of LRRK2 itself, at Ser1292, was used to reflect kinase activity. Increased phosphorylation at this residue has been reported in LRRK2 present in urinary exosomes from both idiopathic as well as G2019S-LRRK2 PD patients ^3,4^. In PBMCs, however, while total LRRK2 expression is readily detectable, phosphorylation at this residue is very weak, unless a signal boost reagent was included. We loaded increasing amounts of PBMC protein extract and probed the membranes for pS1292 and total LRRK2. The inclusion of the Signalboost reagent may alter the linearity of the detection of pS1292-LRRK2 by western immunoblot (Supplemental Fig. 1b) in unknown ways; thus we elected not to use this pharmacodynamics readout in assessing LRRK2 function in PBMCs.

*pS1292-LRRK2 in urine exosomes*. Previous reports have assessed auto-phosphorylation of LRR2 at the Ser1292 site, and found increased levels in urinary exosomes from iPD and G2019S-PD patients ^3,4^. We assessed pS1292-LRRK2 by Western immunoblotting in extracts of urine exosomes. Detection of LRRK2, as well as pS1292-LRRK2, was quite variable in samples across the subject groups, with LRRK2 being un-detectable in many cases (Supplementary Figure 1b).

References.

1	Melachroinou, K. *et al.* Elevated in vitro kinase activity in PBMCs of LRRK2 G2019S carriers: a novel ELISA-based method. *Mov Disord* **in press** (2020).

2	Shi, M. *et al.* Plasma exosomal alpha-synuclein is likely CNS-derived and increased in Parkinson's disease. *Acta Neuropathol* **128**, 639-650, doi:10.1007/s00401-014-1314-y (2014).

3	Fraser, K. B., Moehle, M. S., Alcalay, R. N., West, A. B. & Consortium, L. C. Urinary LRRK2 phosphorylation predicts parkinsonian phenotypes in G2019S LRRK2 carriers. *Neurology* **86**, 994-999, doi:10.1212/WNL.0000000000002436 (2016).

4	Fraser, K. B. *et al.* Ser(P)-1292 LRRK2 in urinary exosomes is elevated in idiopathic Parkinson's disease. *Mov Disord*, doi:10.1002/mds.26686 (2016).
